# Bacterial cGAS-like enzymes produce 2′,3′-cGAMP to activate an ion channel that restricts phage replication

**DOI:** 10.1101/2023.07.24.550367

**Authors:** Uday Tak, Peace Walth, Aaron T. Whiteley

## Abstract

The mammalian innate immune system uses cyclic GMP–AMP synthase (cGAS) to synthesize the cyclic dinucleotide 2′,3′-cGAMP during antiviral and antitumor immune responses. 2′,3′-cGAMP is a nucleotide second messenger that initiates inflammatory signaling by binding to and activating the stimulator of interferon genes (STING) receptor. Bacteria also encode cGAS/DncV-like nucleotidyltransferases (CD-NTases) that produce nucleotide second messengers to initiate antiviral (antiphage) signaling. Bacterial CD-NTases produce a wide range of cyclic oligonucleotides but have not been documented to produce 2′,3′-cGAMP. Here we discovered bacterial CD-NTases that produce 2′,3′-cGAMP to restrict phage replication. Bacterial 2′,3′-cGAMP binds to CD-NTase associated protein 14 (Cap14), a transmembrane protein of unknown function. Using electrophysiology, we show that Cap14 is a chloride-selective ion channel that is activated by 2′,3′-cGAMP binding. Cap14 adopts a modular architecture, with an N-terminal transmembrane domain and a C-terminal nucleotide-binding SAVED domain. Domain-swapping experiments demonstrated the Cap14 transmembrane region could be substituted with a nuclease, thereby generating a biosensor that is selective for 2′,3′-cGAMP. This study reveals that 2′,3′-cGAMP signaling extends beyond metazoa to bacteria. Further, our findings suggest that transmembrane proteins of unknown function in bacterial immune pathways may broadly function as nucleotide-gated ion channels.

## Introduction

Cyclic dinucleotides are important second messengers that play essential roles in cellular physiology and immune signaling throughout the tree of life. In mammalian cells, the enzyme cyclic GMP–AMP synthase (cGAS) produces the cyclic dinucleotide 2′,3′-cyclic GMP–AMP (2′,3′-cGAMP) to initiate an immune response (*1*). 2′,3′-cGAMP is produced when cGAS is directly activated by binding cytosolic double-stranded DNA, which becomes available during infection and cellular stress (*1*). 2′,3′-cGAMP diffuses throughout the cell and binds to Stimulator of Interferon Genes (STING), which activates downstream inflammatory pathways and type I interferon signaling (*1, 2*). In this way, cGAS acts as a sensor and the cGAS-STING pathway is crucial for immune responses to viruses, intracellular pathogens, and cancer (*1, 2*).

Bacteria encode pathways related to metazoan cGAS-STING called cyclic oligonucleotide-based antiphage signaling systems (CBASS), which protect against phage infection (*3*–*7*). CBASS operons encode structural homologs of cGAS called cGAS/DncV-like nucleotidyltransferases (CD-NTases) that produce cyclic di- and oligonucleotides during phage infection (*3, 4, 8*–*10*). The cyclic oligonucleotide binds to and activates a cognate CD-NTase-associated protein (Cap), including STING-like effectors, which activates the protein (*3, 4, 6, 11*–*16*). In general, Cap effectors initiate programmed cell death during phage infection to prevent the virus from completing its replication cycle, thus allowing the bacterial population to survive (*17*). CBASS pathways signal via a range of cyclic nucleotides unique to bacteria, including cyclic trinucleotides, 3′,3′-cGAMP and 3′,2′-cGAMP, however, the mammalian 2′,3′-cGAMP molecule has not yet been found in bacteria (*3, 12, 16, 18*). Given the specificity of human cGAS and potent signaling initiated by 2′,3′-cGAMP in mammalian cells, we sought to identify whether bacteria also encode 2′,3′-cGAMP signaling pathways.

## Results

### Bacterial CdnB enzymes produce 2′,3′-cGAMP

We searched for 2′,3′-cGAMP production in bacteria by screening diverse CD-NTases for production of 2′,3′-cGAMP *in vitro*. Previously, we incubated recombinant CD-NTases with α^32^P-radiolabelled ATP, CTP, GTP and UTP and screened for cyclic oligonucleotide production by thin layer chromatography (**Fig. 1A**) (*3*). Here we extended this analysis by digesting these products with P1 endonuclease, an enzyme that exclusively hydrolyzes 3′–5′ phosphodiester-linked nucleotides. Cyclic oligonucleotides with only 3′–5′ phosphodiester linkages are completely cleaved by this enzyme. However, 2′,3′-cGAMP contains one 2′–5′ and one 3′–5′ phosphodiester linkage, therefore this molecule is only partially digested, and the remaining 2′–5′-linkage is preserved. Using this analysis, we found at least four CD-NTase products that harbored non-3′–5′ linkages **(Fig. 1B)**.

**Figure 1.**
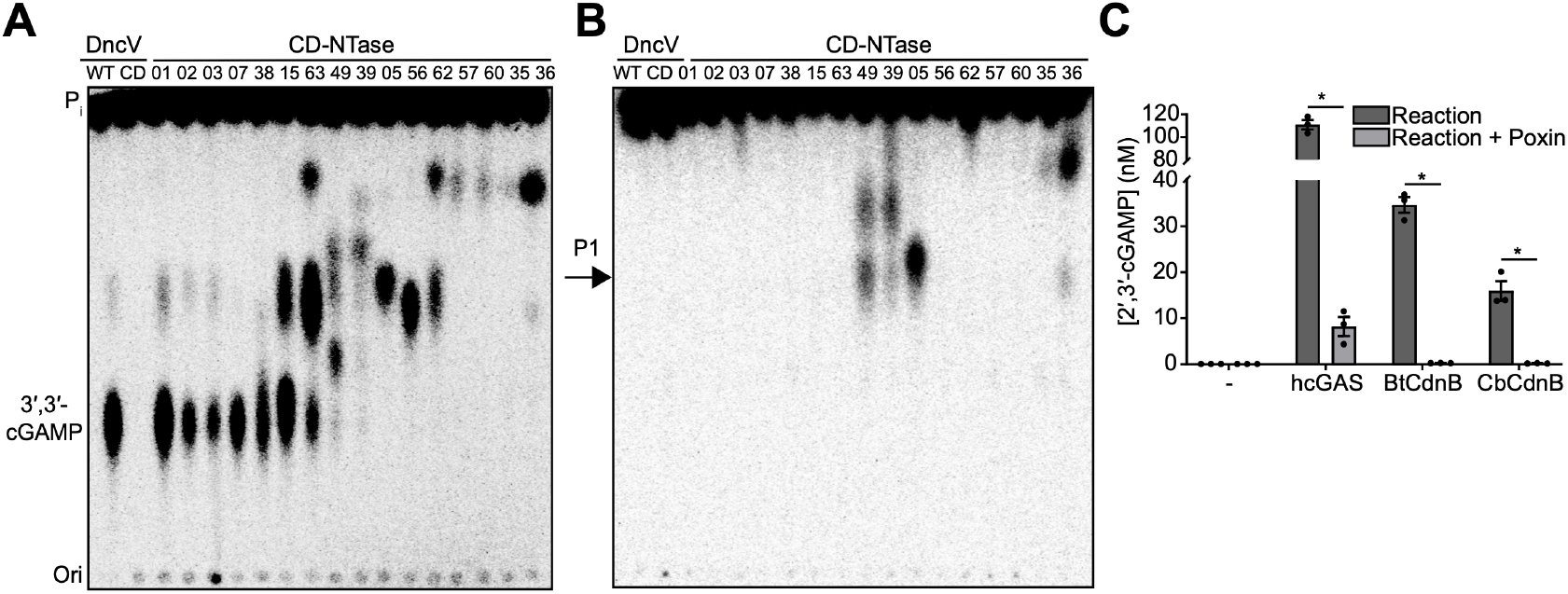
Bacterial cGAS-like enzymes produce 2′,3′-cGAMP. **(A)** PEI-cellulose TLC analysis of enzyme reaction products produced when the indicated CD-NTases were incubated with α^32^P-radiolabelled ATP, CTP, GTP and UTP. *Vibrio cholerae* DncV was used to produce 3′,3′-cGAMP in lane one. Catalytically dead (CD) DncV^DID79AIA^; inorganic phosphate (Pi); origin (Ori.). Images are representative of n = 2 biological replicates. **(B)** Analysis as in (a) after treatment with nuclease P1 (P1), which cleaves 3′–5′ phosphodiester bonds. Spots remaining after nuclease P1 treatment contain non-3′–5′ linked ^32^P, such as 2′–5′ linkages. **(C)** ELISA quantification of 2′,3′-cGAMP from enzyme reaction products produced when the indicated enzyme was incubated with ATP and GTP. Where indicated, reaction products were further treated with VacV Poxin, a viral enzyme that specifically degrades 2′,3′-cGAMP ((19,20). Data are mean ± standard error of the mean (SEM) for technical replicates and representative of n = 2 biological replicates.

We selected CD-NTase005 from *Desulfotomaculum alkaliphilum* (Da) for further analysis and deconvolution of the substrate specificity revealed that this enzyme used ATP and GTP as substrates **(Fig. S1A)**, consistent with production of 2′,3′-cGAMP. Sequence analysis of CD-NTase005 places this enzyme in Clade B02 of the CD-NTase phylogeny **(Fig S1B)**. We renamed this enzyme DaCdnB and analyzed additional CdnB homologs by thin layer chromatography. CdnB homologues from *Bacillus thuringiensis* (BtCdnB) and *Clostridium botulinum* (CbCdnB) **(Fig. S1C)** produced CDN products similar to DaCdnB, which appeared to be linear pppG(2′,5′)pA. Linear pppG(2′,5′)pA is the reaction intermediate produced by cGAS prior to final cyclization into 2′,3′-cGAMP. We speculated that production of the linear intermediate was due to non-optimal CdnB reaction conditions. To overcome this issue, we included manganese (a known activator of nucleotidyltransferases) in our reactions and found that BtCdnB robustly produced a cyclic dinucleotide that migrated similarly to 2′,3′-cGAMP **(Fig S1C)**.

To test the hypothesis that CdnB produced 2′,3′-cGAMP, we evaluated the reaction products of BtCdnB, CbCdnB, and human cGAS using a 2′,3′-cGAMP-specific ELISA. Both BtCdnB and CbCdnB produced high levels of 2′,3′-cGAMP similar to activated human cGAS **(Fig 1C)**. Measurement of 2′,3′-cGAMP using this ELISA was highly specific as we did not detect any cross-reactivity using synthetic 3′,2′, 3′,3′-cGAMP, linear pppG(2′,5′)pA, cyclic di-AMP, or cyclic di-GMP, and negligible signal for 2′,2′-cGAMP **(Fig. S1E)**. Further, the 2′,3′-cGAMP produced by BtCdnB and CbCdnB was destroyed by recombinant VacV poxin protein **(Fig. 1C)**, which exclusively hydrolyzes 2′,3′-cGAMP (*19, 20*). Taken together, these experiments reveal that CdnB enzymes from *B. thuringiensis* and *C. botulinum* produce 2′,3′-cGAMP.

### CdnB signaling restricts bacteriophage replication

Analysis of the genomic context of *cdnB* showed that these genes are typically encoded in type I CBASS systems: two-gene operons that consist of a CD-NTase and a putative nucleotide-activated effector protein (*5, 7*). The putative effector frequently adjacent to *cdnB* is CD-NTase-associated protein 14 (*cap14*), a gene of unknown function (Fig. 2A) (*7, 13*). We selected a CBASS system encoding *cdnB-cap14* from *Bacillus thuringiensis* 407 (BtCBASS) and expressed these genes under their endogenous promoter in *Bacillus subtilis*. BtCBASS provided over 10^4^-fold protection against phages SPP1 and phiB002 **(Fig. 2B, C, D)** but did not protect against phages SPβ or Φ29 **(Fig. Fig. S2A, B)**. Phage defense was dependent on the BtCdnB active site residues, which are required for cyclic oligonucleotide production **(Fig. 2B, C, D)**.

**Figure 2.**
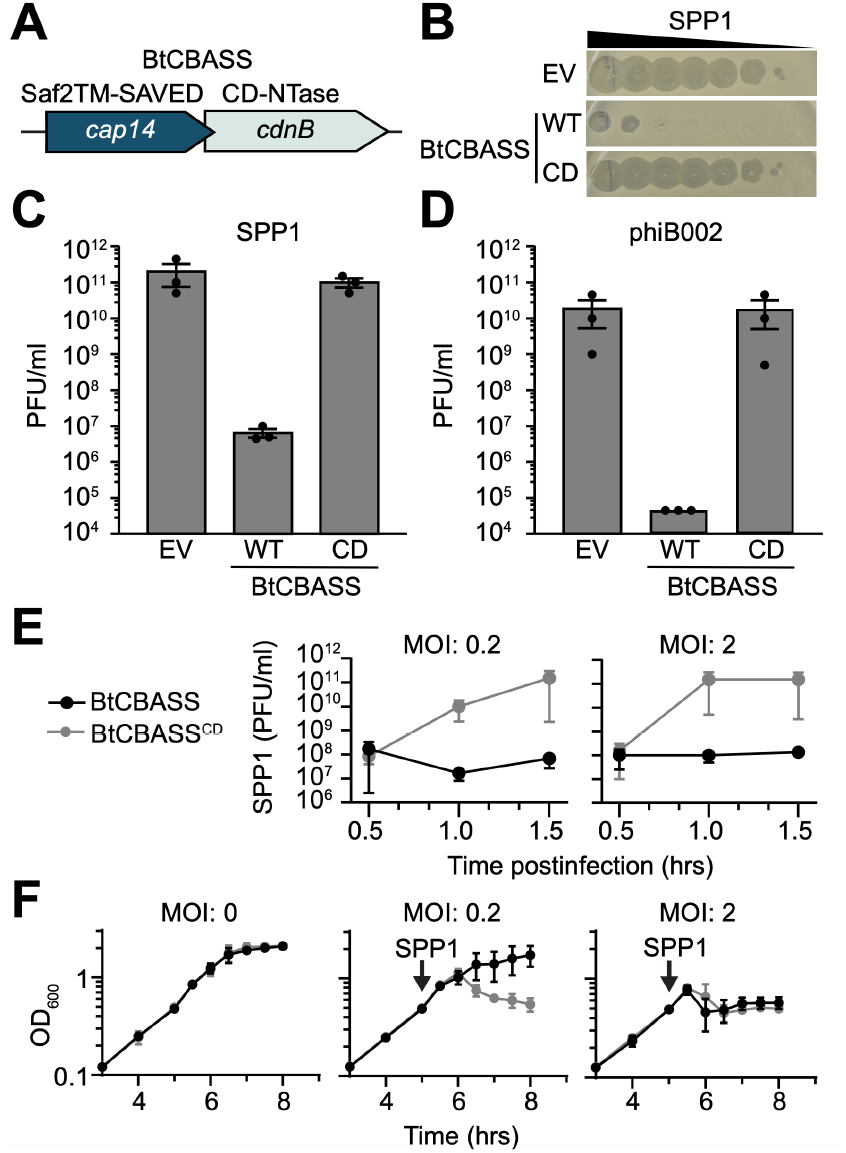
BtCBASS protects bacteria against phage. **(A)** BtCdnB is encoded by *Bacillus thuringiensis* 407 in a type I CBASS system (BtCBASS). These systems encode a CD-NTase and a predicted effector protein that is typically activated by nucleotide binding. The predicted effector protein for BtCBASS is CD-NTase-associated protein 14 (*cap14*), which encodes an N-terminal Saf-2TM and a C-terminal SAVED domain. **(B)** Efficiency of plating of phage SPP1 infecting *Bacillus subtilis* expressing BtCBASS, catalytically dead BtCBASS (CD) encoding CdnB^DID79AIA^, or an empty vector (EV). Data are representative images from n = 3 biological replicates. **(C and D)** Efficiency of plating of phage SPP1 or phiB002 (isolated for this study) infecting *B. subtilis* expressing the indicated genotype. Data represent mean ± SEM for n = 3 biological replicates. **(E)** PFU/ml of phage SPP1 during liquid culture infection of *B. subtilis* containing BtCBASS or the CD strain. Shown are the plaque-forming units per milliliter (PFU/ml) at the indicated time points after infection of mid-log phase (OD = 0.4) bacteria at a multiplicity of infection of 0.2 or 2. **(F)** Growth curve of *B. subtilis* expressing the indicated genotype. Arrows indicate infection with phage SPP1 at the indicated multiplicity of infection (MOI). Data represents the mean ± SEM of n = 3 biological replicates.

BtCBASS activity had no effect on bacterial growth in the absence of phage and completely prevented a collapse in culture OD_600_ when phage was added at a multiplicity of infection (MOI) of 0.2 **(Fig. 2F)**. The decrease in culture OD_600_ was due to phage replication as strains expressing catalytically inactive BtCBASS supported a 1000-fold increase in plaque forming units over wild-type BtCBASS **(Fig. 2E)**. When the MOI was increased to 2, such that every bacterium was infected, the OD_600_ of cultures expressing wild-type and catalytically inactive BtCBASS both collapsed. However, bacteria expressing wild-type BtCBASS still failed to support viral replication in this condition **(Fig. 2E, F)**. The precipitous drop in culture OD_600_ when all BtCBASS-expressing bacteria are infected suggests that activation of BtCBASS results in an abortive infection phenotype. The similar kinetics of OD_600_ decrease in cultures expressing active and inactive BtCBASS is consistent with other reports demonstrating that CBASS systems are activated late in infection (*4, 21*). These data demonstrate that BtCdnB-mediated nucleotide second-messenger synthesis is required for BtCBASS function and phage resistance.

### Cap14 is a 2′,3′-cGAMP-activated ion channel

To determine the mechanism for BtCdnB-mediated phage resistance, we focused on the putative nucleotide receptor BtCap14, the second protein encoded by BtCBASS. Cap14 proteins were previously predicted to have an N-terminal transmembrane 2TM module named Saf-2TM (SAVED-fused 2TM) and a C-terminal SAVED domain (*7*). SAVED domains are commonly found in effector proteins of CBASS systems where they bind CD-NTase-derived nucleotides to coordinate activation of a fused protein domain. SAVED domains fused to nucleases and TIR domains have been previously investigated (*12, 14, 16*), but SAVED domains fused to predicted transmembrane domains have not been investigated biochemically. We expressed and purified full-length BtCap14 in detergent micelles **(Fig. S3A)** and measured cGAMP binding. BtCap14 bound 2′,3′-cGAMP with a K_D_ of 11.3 ±0.83 nanomolar but did not appreciably bind other cGAMP isoforms **(Fig. 3A)**. These data confirm that Cap14 is indeed a nucleotide receptor and provide additional evidence that BtCBASS signals via 2′,3′-cGAMP.

**Figure 3.**
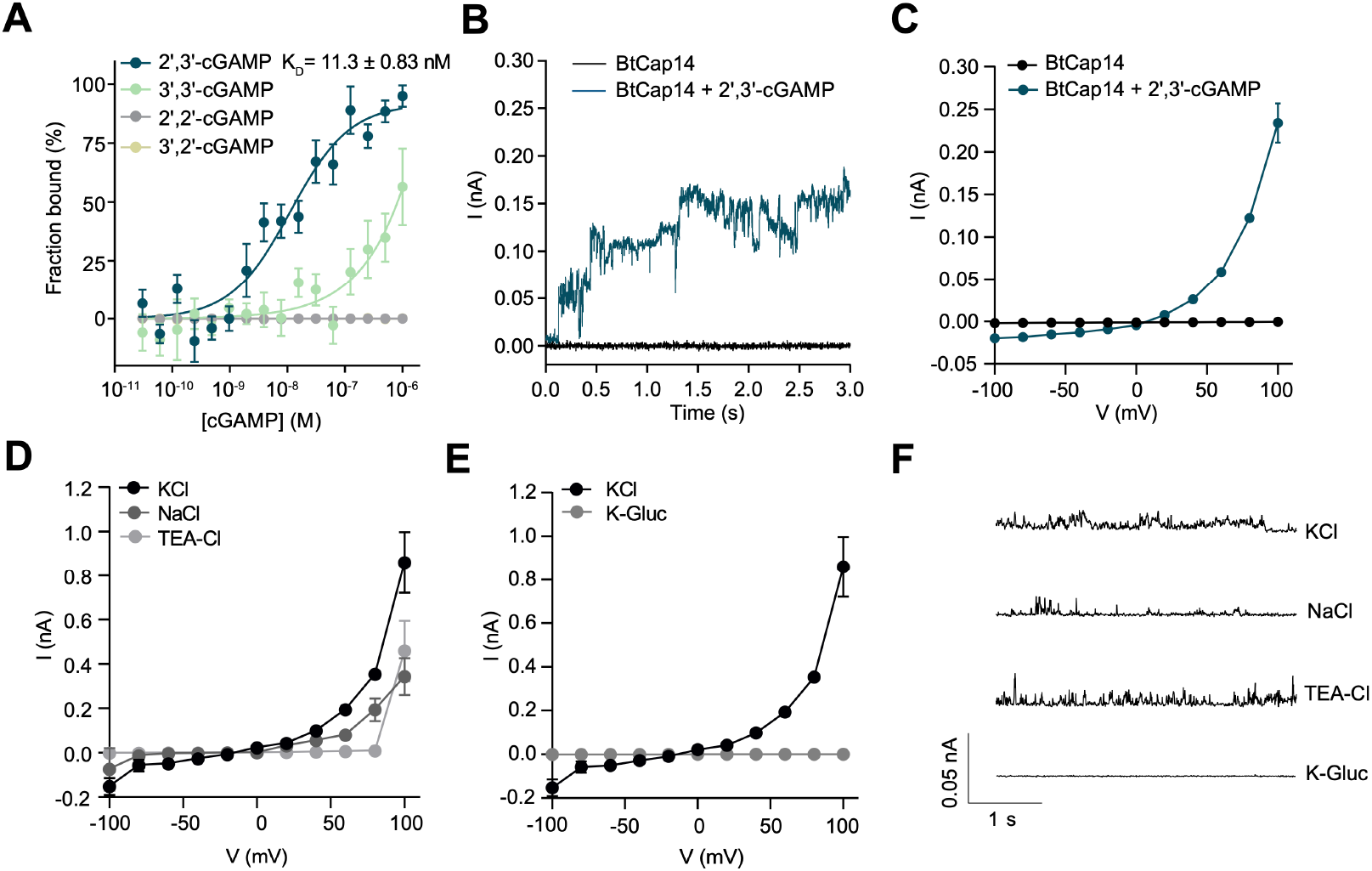
BtCap14 is a 2′,3′-cGAMP-activated ion channel. **(A)** Microscale thermophoresis analyzing binding of BtCap14_6xhis_ to the indicated cGAMP isoforms. Data are n = 3 biological replicates for 2′,3′-cGAMP, 3′,3′-cGAMP, 2′,2′-cGAMP, and n=2 for 3′,2′-cGAMP. **(B)** Voltage clamp electrophysiology of BtCap14_6xhis_ in planar lipid bilayers. Shown is a representative current trace of BtCap14 with or without 1 µM 2′,3′-cGAMP added to the *cis* side of the bilayer. Bilayers were formed with 1,2-diphytanoyl-sn-glycero-3-phosphocholine (DphpC) lipid bilayers in a symmetric solution of 10 mM HEPES pH 7.2, 1 M KCl on *cis* and *trans* sides of the bilayer, and recordings were performed at a holding voltage of +40 mV. Data are representative of n ≥ 3 biological replicates. **(C)** Current / voltage relationship (I/V) of BtCap14 in the apo state (without cGAMP) and in the presence of 1 µM 2′,3′-cGAMP added to the *cis* side of the bilayer. Data is the mean ± standard deviation (SD) for a representative experiment from n ≥ 3 biological replicates. **(D)** I/V relationship of BtCap14 + 1 µM 2′,3′-cGAMP in 1 M of the indicated salt replacing KCl as the recording solution under symmetric conditions. Data is the mean ± SD for a representative experiment from n ≥ 3 biological replicates. **(E)** I/V relationship of BtCap14 + 1 µM 2′,3′-cGAMP in symmetric 10 mM HEPES pH 7.2 + 1 M KCl or 1 M K-Gluconate as the recording solution. Data is the mean ± SD for a representative experiment from n ≥ 3 biological replicates. **(F)** Single channel recording of BtCap14 + 1 µM 2′,3′-cGAMP at +100 mV in 10 mM HEPES pH 7.2 + 1 M of the indicated salt in symmetric conditions. Data are representative of n ≥ 3 biological replicates.

Analysis of the BtCap14 amino acid sequence identified three putative transmembrane helices in the Saf-2TM domain **(Fig. S3B)**. However, a model of the predicted structure for the BtCap14 monomer did not show Saf-2TM as uniformly hydrophobic but rather as having opposing hydrophobic and hydrophilic faces, similar to pore-forming subunits of multimeric ion channels **(Fig. S3C, D)**. Gel filtration revealed that BtCap14 eluted as an oligomer **(Fig. S3E)** and structure predictions were used to produce models for a tetramer **(Fig. S3F)**. The architecture of the tetramer resembled an ion channel in which the hydrophobic faces of each monomer pointed outward into the membrane and the hydrophilic faces pointed inwards creating a putative conduit for ion transport **(Fig. S3F)**.

To test whether BtCap14 displays ion channel activity we reconstituted the detergent-solubilized complex in artificial lipid bilayers and directly measured ion flux across the membrane using voltage-clamp electrophysiology **(Fig. S4A)** (*22*–*24*). In this assay, the voltage is held constant creating a driving force, and ion flux is measured by the resulting change in current. BtCap14 alone did not alter the current trace from baseline, however, addition of 2′,3′-cGAMP to the *cis* chamber resulted in stepwise increases in the current trace **(Fig 3B)**. This pattern is indicative of ion flux across the membrane through the opening of BtCap14 pores and is a hallmark of channel-forming proteins. The channel activity of 2′,3′-cGAMP-activated BtCap14 was sensitive to the compound gadolinium chloride (Gd^3+^), a known inhibitor of ion channels **(Fig. S4B)**.

The current/voltage analysis of BtCap14 in a symmetrical KCl solution confirmed that observed channel activity is strictly dependent on 2′,3′-cGAMP **(Fig 3C)**. Unexpectedly, 2′,3′-cGAMP-activated BtCap14 displayed channel activity only at positive holding potentials **(Fig 3C, S4C, D)**. This behavior is indicative of ion selectivity and/or directionality of ion flux (*23*). To define the ion(s) capable of transiting the BtCap14 channel, we substituted the recording solution with varying cations and anions then measured activity at +100 mV holding potential. BtCap14 current/voltage profiles and single channel activity were unaffected when KCl was replaced with NaCl or tetraethylammonium chloride (TEA-Cl). TEA^+^ (∼ 3.85 Å ionic radius) is much larger than K^+^ (∼ 1.3 Å ionic radius) or Na^+^ (∼ 1 Å ionic radius) and is often impermeable to known cation channels (*25*) **(Fig. 3D, F)**. Therefore, we hypothesized that BtCap14 may be an anion transporter. In line with that hypothesis, substitution of KCl with potassium gluconate (K-Gluc) resulted in a complete lack of channel activity **(Fig. 3E, F)**. In the presence of 2′,3′-cGAMP, buffer exchange of K-Gluc with KCl on the *cis*-side of BtCap14 bilayers resulted in a recoverable current fluctuation **(Fig. S4E)**. These experiments suggest that active BtCap14 transports Cl^-^ from the *cis* to *trans* chambers.

BtCap14 undoubtably inserts into the artificial lipid bilayer in multiple orientations, however, 2′,3′-cGAMP was only added to the *cis* chamber and does not cross membranes. Therefore, only BtCap14 with SAVED domains oriented on the *cis* side of the bilayer can be activated. In this way, the *cis* chamber mimics the bacterial cytosol, where the SAVED domain would be accessible to 2′,3′-cGAMP produced by BtCdnB in response to phage infection. As our data show Cl^-^ transport from *cis* to *trans*, our findings suggest that BtCap14 is a nucleotide-gated ion channel capable of effluxing Cl^-^ ions from the cytosol to the extracellular space.

### Generation of a 2′,3′-cGAMP biosensor

2′,3′-cGAMP is a crucial signaling molecule in mammalian cells. Indeed, cGAS-STING signaling plays a major role in responsiveness to cancer immunotherapy (*26*–*28*). However, a key challenge in the study of 2′,3′-cGAMP is accurate, sensitive, and rapid quantification of nucleotide in complex mixtures, such as cell lysates. We reasoned that our findings on the specificity and high affinity of Cap14 for 2′,3′-cGAMP might help overcome this challenge if the nucleotide binding domain of Cap14 could be used to construct a biosensor. Native Cap14 is not well suited as a biosensor because electrophysiology is technically challenging and susceptible to bilayer-disrupting contaminants. However, if the nucleotide-binding “input” domain of Cap14 could be appended to an enzymatic “output” domain, then a chimeric protein could link 2′,3′-cGAMP concentration with enzymatic activity. This chimeric approach appeared possible because the predicted nucleotide-binding domain of Cap14 is a C-terminal SAVED domain, a conserved structure encoded naturally as a fusion with a wide range of N-terminal enzymatic effector domains (*7*).

We constructed a series of chimeric proteins that fused the C-terminal SAVED domain of Cap14 homologs with N-terminal HNH nuclease domains of Cap5 homologs and screened these proteins for nuclease activity in the presence or absence of 2′,3′-cGAMP. Most of these proteins displayed poor expression or solubility, however, a fusion of the HNH nuclease domain from *Geobacillus* Cap5 to the SAVED domain of *C. botulinum* Cap14 **(Fig 4A)** was readily expressed and purified from *E. coli* **(Fig S5A)**. This protein was named Chimera10. Purified Chimera10 did not degrade a DNA substrate in the absence of nucleotide, however, when incubated with 2′,3′-cGAMP we observed DNA degradation, indicative of nuclease activity **(Fig. 4B)**. No other cGAMP isomers or common cyclic dinucleotides were capable of activating DNA cleavage by Chimera10 **(Fig. 4B)**. Using a fluorescent DNA-cleavage reporter assay, we determined that Chimera10 could be activated by 10 nM 2′,3′-cGAMP, which is consistent with the native ligand affinity found in the related BtCap14 **(Fig. 4C)**. These experiments provide proof-of-concept that SAVED domains are modular and can be functionally exchanged between effector types to program ligand-specific activation.

**Figure 4.**
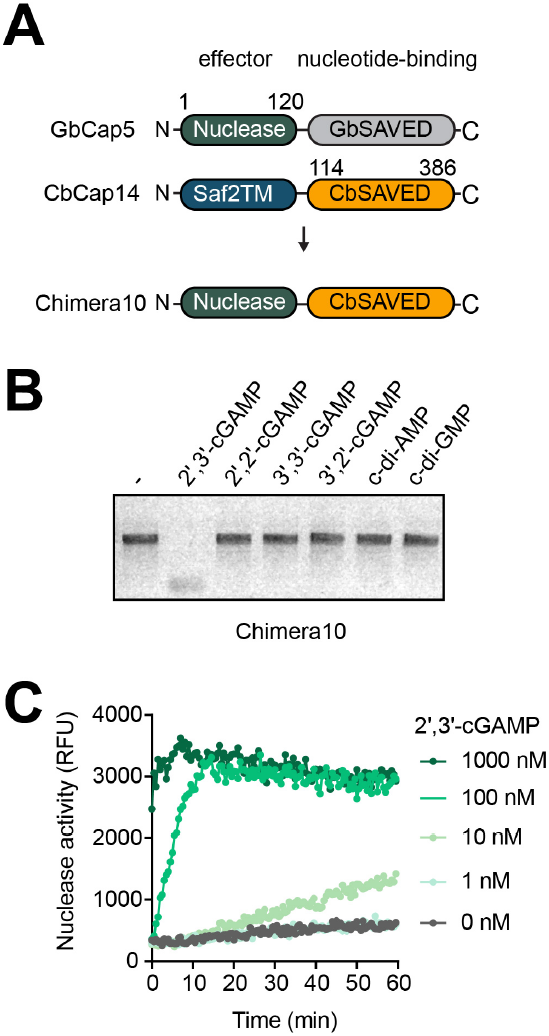
Domain-swapping of CBASS effectors enables engineering of a 2′,3′-cGAMP biosensor. Receptor-swapping strategy for engineering a 2′,3′-cGAMP-activated biosensor. GbCap5 (*Geobacillus*) contains an N-terminal HNH endonuclease with a C-terminal SAVED domain that responds to 3′,2′-cGAMP and 2′,3′-cGAMP at 1 µM (data not shown). The SAVED domain from *Clostridium botulinum* (CbCap14) was swapped with the GbSAVED domain to construct the chimeric nuclease effector: Chimera10. Numbers represent amino acid positions. DNA degradation assay. Linear DNA was incubated with Chimera10 and 1 µM of the indicated cyclic dinucleotide for 30 minutes at 37 °C. Reactions were separated by agarose gel electrophoresis and visualized. Data are representative of n = 3 biological replicates. **(C)** Fluorescence-based DNase activity assay of Chimera10 activity. Cleavage of the substrate results in fluorescence was measured as relative fluorescence units (RFU), which was monitored over time. A representative experiment is shown from n = 2 biological replicates.

## Discussion

In this study, we identified a subset of bacterial CdnB enzymes that produce 2′,3′-cGAMP to activate Cap14: a novel nucleotide-gated ion channel, to restrict phage replication. Previously, 2′,3′-cGAMP has only been identified in metazoans where it is produced by cGAS and allosterically activates STING. A unique quality of 2′,3′-cGAMP signaling in mammals is that extracellular 2′,3′-cGAMP can be imported by mammalian cells to activate STING in paracrine (*29*–*33*). As STING binds 2′,3′-cGAMP with high affinity (K_D_ of ∼4 nM) (*34*) and leads to potent type I interferon induction, even minute amounts of extracellular 2′,3′-cGAMP could have impacts on immune signaling. This raises the intriguing possibility that bacteria producing 2′,3′-cGAMP may alter signaling within nearby mammalian cells. Our analysis identified CdnB-encoding bacteria associated with metazoan hosts, including *Bacillus, Clostridium, Enterococcus*, and *Staphylococcus* species (Fig S1B). Further, some of these homologs are encoded by bacteria that are members of the human microbiome. It is therefore plausible that bacterial 2′,3′-cGAMP could be released following CBASS-mediated bacteriolysis and activate mammalian STING. This hypothesis is especially interesting because STING signaling has wide-reaching implications for host-pathogen interactions, antitumor immunity, and the efficacy of host-directed cancer immunotherapy (*2, 23, 25, 32*). Given there are multiple correlations between microbiome constituents and disease/treatment of diseases (*35*– *38*), perhaps some of these may be explained by bacterial 2′,3′-cGAMP production. Our findings are limited to the CdnB02 subclade of CD-NTases, however there are likely additional 2′,3′-cGAMP synthases that are yet to be discovered. Further exploration of bacterially-derived 2′,3′-cGAMP may yield new mechanistic explanations linking microbiota and human health and immunity. Consequently, these findings may inform new strategies for designing therapeutics for host-directed immunotherapy in cancer and infectious disease.

Exogenous delivery of 2′,3′-cGAMP is also a promising strategy to activate STING-mediated signaling for host-directed immunotherapy in infectious disease and cancer (*27*). These approaches rely on liposome or nanoparticle-based delivery (*39*), development of small-molecule STING agonists (*40, 41*), or engineering of human cGAS to constitutively produce 2′,3′-cGAMP (*42*). CdnB enzymes provide a new method to introduce 2′,3′-cGAMP in a therapeutic setting. These enzymes are constitutively active *in vitro* and could be exploited further as fusion proteins to potentiate existing therapies.

In addition, our study led to development of a new biosensor for 2′,3′-cGAMP by investigating the CBASS effector Cap14. Sequence analysis of SAVED-domain-containing proteins predicted that this domain is a module that can mediate activation of a wide-variety of effector domains involved in biological conflict. Here, we demonstrate that indeed SAVED domains can be exchanged to reprogram CBASS effectors. This quality of SAVED is likely key to the evolution of CBASS systems through generating effector and nucleotide second messenger diversity, which may be an evolutionary advantage for bacteria in their conflict with phage and ecological niche (*7, 43*). In this study, we used SAVED reshuffling to construct Chimera10 and link 2′,3′-cGAMP with nuclease activity. Detection of 2′,3′-cGAMP is currently limited to STING-based sensors (*44*), RNA-based probes (*45*), and endpoint assays such as mass-spectrometry (*46*). A distinct advantage of Chimera10 is exquisite selectivity for 2′,3′-cGAMP over other isomers and alternative cyclic dinucleotides. This specificity and modularity of SAVED-containing effectors offers a new platform for development of 2′,3′-cGAMP-activated biotechnology.

The Cap14 effector in the BtCBASS system functions as a 2′,3′-cGAMP-gated ion channel. Transmembrane effectors are highly abundant in bacterial antiphage systems (*47*–*52*). The CBASS Cap15 and Cap14 effectors, and Pycsar pycTM effectors were observed to deform membranes during phage infection of *E. coli* but the biochemical basis was not elucidated (*13, 53*). This phenomenon could occur via effector association with existing membrane components to induce membrane deformation or through direct pore-formation by the effector itself. Our biochemical data supports the latter model for Cap14, although other TM effectors may function differently. Although BtCap14 is selective for Cl^-^, it is possible that other Cap14 effectors possess different properties that are dictated by variability in the Saf-2TM domain. Intriguingly, Cap14’s ion selectivity distinguishes it from other pore-forming immune effectors such as gasdermins, which insert into the membrane from water-soluble precursors to form large-diameter nonselective pores (*54, 55*). It remains to be determined exactly how Cl^-^ efflux protects bacteria against phage, although our data showing large changes in culture OD_600_ suggest bacteriolysis and abortive infection. It will be of particular interest to understand how phages overcome channel-mediated immunity as they do for other bacterial immune systems (*56*). Pore-formation is a conserved feature of many programmed cell death pathways in animals, plants, fungi, and bacteria (*25, 54, 55, 57*–*61*). Our findings expand this paradigm and suggest that direct ion channel activity executes antiviral activity in bacteria. We predict that many other immune effectors, from bacteria and other phyla, may act as novel ion channels.

## Acknowledgements

The authors would like to thank Dr. Kelly Doran (University of Colorado Anschutz Medical Campus) for use of their Nanion Orbit Mini planar lipid bilayer device; Dr. Alexander Horswill (University of Colorado Anschutz Medical Campus) for providing *E. coli* C43; Dr. Daniel Kearns (Indiana University) for phage SPP1; Dr. Deborah Wuttke (University of Colorado Boulder) for use of their FPLC; Dr. David Rudner (Harvard University) for *Bacillus subtilis* PY79 strain; the CU Boulder Department of Biochemistry Shared Instruments Pool core facility (RRID:SCR_018986), Dr. Annette Erbse, and its staff, including use of their MST instrument (NIH: S10OD21603); Hannah Arthur for assistance with isolation of wild *B. subtilis* phages; Dr. Joel Kralj and Dr. Christian Meier for helpful discussions and the use of their plate reader; and members of the Aaron Whiteley lab for advice and helpful discussion.

## Author contributions

Conceptualization: U.T, and A.T.W.; Methodology, U.T. and A.T.W, Investigation., U.T, P.H.W, and A.T.W, Writing – Original Draft, U.T, and A.T.W.; Visualization, U.T and A.T.W.; Supervision, A.T.W.; Funding Acquisition, U.T and A.T.W.

## Data and materials availability

All data and materials generated in this study are available upon request.

## Materials and Methods

### Cloning and Plasmid Construction

All inserts encoding for the indicated proteins or operons were amplified using Q5 polymerase, visualized by SYBR-SAFE-stained agarose gel electrophoresis, and extracted using the Qiagen gel extraction kit. pLOCO2 vectors created for *Bacillus subtilis* expression used inserts containing SbfI/ NotI cut sites, pET vectors for 6xhis-SUMO fusion proteins were cloned with BamHI/ NotI sites, pET vectors for C-terminal-6x-his tag proteins were cloned with NdeI/ BamHI sites. Plasmids were assembled via Gibson Assembly (New England BioLabs 2X Master mix) (*62*) using the manufacturers protocol for 1 hour at 50°C. The entire reaction was transformed into chemically competent *E. coli* Omnipir (*9*) using heat shock, outgrown in S.O.C media for 1 hour at 37°C at 200 RPM, followed by selection on Lysogeny-broth (LB) agar supplemented with the required antibiotic + 1% glucose in an incubator at 37°C. Clones were grown in LB + 1% glucose + antibiotic overnight and stored in 15% glycerol at -80°C. Clones were validated using colony PCR and plasmids were isolated using the Qiagen Miniprep kit. Plasmid inserts were validated using Sanger Sequencing (Quintara Biosciences or Genewiz/ Azenta) or using Plasmidsaurus for whole plasmid sequencing. Map construction, primer design, and sequencing analysis were all performed using Geneious Prime.

### Thin layer chromatography and P1 endonuclease treatment

Thin layer chromatography and radioactive CD-NTase enzymatic assays were performed as described previously (*3*).

### Bioinformatic identification of CD-NTases

All CD-NTase numbers and clade assignments are based on the original description from Whiteley et al., *Nature* 2019 (*3*). The amino acid sequence for CD-NTase005 was used as a BLAST seed in Geneious Prime to identify homologues in bacteria. The E-value was used to curate hits, and then relevant hits were aligned using MUSCLE to confirm similarity. These results were then used to make a phylogram in Geneious Prime. Species deemed relevant to our study, human health, or agriculture were included for brevity. Redundant species were excluded for brevity.

### Recombinant Protein Expression and Purification

MBP-CD-NTase constructs used in Fig.1a, b and Fig. S1a, c, were purified as described previously (*3*). For purification of MBP-human cGAS; the plasmid encoding MBP-TEV-hcGAS was transformed into BL21(DE3) and selected on LB-agar supplemented with Cm20 + 1% glucose in an incubator overnight at 37°C. A single colony was grown in 50 ml of LB Cm20 at 37°C until the optical density at 600 nm (OD_600_) reached 0.5, and then 0.2% arabinose was added to induce expression and the culture was moved to 20°C for 24 hours. Cells were harvested and lysed as described above and were run over 5 ml of Amylose affinity resin (New England Biolabs), washed with 100 ml of 20 mM sodium phosphate pH 7.2, 500 mM NaCl, 1 mM DTT, and eluted in the same buffer + 30 mM maltose monohydrate. Samples were concentrated and stored in 50% glycerol in aliquots at -20°C until needed.

CD-NTase005 and its homologues were expressed as 6x-His-SUMO fusion proteins in *E. coli* BL21 Rosetta cells. Plasmids were transformed into Rosetta using heat shock and plated on LB-agar supplemented with carbenicillin 100 µg/ml (Carb100) + chloramphenicol 20µg/ml (Cm20) + 1% glucose and grown in an overnight at 37°C. A single colony was then used to inoculate a 50 ml starter culture of LB Carb100/Cm20 + 1% glucose which was grown overnight at 37°C. The next morning, 10 ml of the overnight culture was used to inoculate 1L of ZYP5052 Studier’s Autoinduction media in 2.5L Thompson flasks (500 ml / flask). Cultures were grown at 37°C for 8 hours and then switched to 20°C for 24 hours. Cells were harvested by centrifugation at 4000 x *g* and resuspended in Buffer 1: 50 mM sodium phosphate pH 7.2, 500 mM NaCl, 10 mM imidazole, 10% glycerol, 1 mM DTT + 1 mg/ml lysozyme. Cells were sonicated on ice at an amplitude of 70, for 30 seconds on / off for 10 minutes total. Sonicated lysates were centrifuged at 14,000 x *g* for 1 hour at 4°C to pellet debris, and the supernatant was run over 2 ml of Ni-NTa resin (Thermo Fisher Scientific) equilibrated with Buffer 1. The resin was washed with 100 ml of Buffer 2: 20 mM sodium phosphate pH 7.2, 500 mM NaCl, 20 mM imidazole, 1 mM DTT to remove nonspecific components bound to the resin. Proteins were eluted using Buffer 3: 20 mM sodium phosphate pH 7.2, 500 mM NaCl, 500 mM imidazole, 1 mM DTT. The 6xhis-SUMO tag was cleaved using 6xhis-ULP1 (produced in-house) via dialysis overnight at 4°C against 4L of Buffer 4: 20 mM sodium phosphate pH 7.2, 500 mM NaCl, 1 mM DTT using a 10 kDa MWCO dialysis membrane. The cleaved sample was run over 2 ml Ni-NTA to capture cleaved 6xhis-SUMO tag and uncleaved protein and the flowthrough containing the recombinant tag-free CD-NTase was collected. CD-NTases were then concentrated via 3kDa ultracentrifugation filter at 4000 x *g* at 4°C and stored in 50% glycerol at -20°C until needed. Protein purity was assessed via SDS-PAGE followed by Colloidal Coomassie staining. 6xhis-SUMO-VacV Poxin was expressed in BL21 (DE3), and purified and stored identically as described above, but was left as a fusion protein since this does not affect activity. The same protocol was used for expression, purification, and storage of GbCap5 and Chimera 10, with the exception that BL21 (DE3) was used as the expression host. Samples were concentrated and buffer exchanged into reaction buffer: 10 mM Tris-HCl pH 7.4, 25 mM KCl, 20 mM MgCl_2_, 1 mM DTT using a centrifugation column with a 3 kDa MWCO (Pall Corporation) prior to use and/or storage. It should be noted that Chimera10 tends to be unstable after 24 hours, and thus experiments were performed immediately after purification.

For expression of recombinant BtCap14; The pAW1517 plasmid was transformed into *E. coli* C43 competent cells (A kind gift from Dr. Alexander Horswill, University of Colorado Anschutz Medical Campus) via heat shock and was plated on LB agar + 100 µg/ml carbenicillin with 1% glucose and grown overnight at 37°C. The following day a single colony was used to inoculate 50 ml of LB + 1% glucose + 100 µg/ml carbenicillin, and the culture was grown overnight for 16 hours at 37°C at 200 RPM. The 50 ml culture was used to inoculate 4L of ZYP5052 autoinduction medium + 1X Vitamin Mix (Teknova) + 100 µg/ml carbenicillin (Note: the ZY base medium was modified to include 20 g tryptone, 10 g yeast extract, 5 g NaCl for 1L). Cultures were split into 2.5L Thompson flasks (500 ml / flask) and were grown shaking for 8 hours at 37°C, 200 RPM and then lowered to 20°C and grown for an additional 24 hours at 200 RPM. Cultures were harvested at 4000 x *g* and resuspended in 40 ml lysis buffer per liter of culture (50 mM sodium phosphate, 500 mM NaCl, 10% glycerol, pH 7.0 + 1 mM PMSF, 2 µl benzonase, + 1 complete EDTA-free protease inhibitor tablet (Thermo Fisher Scientific) + 1 mg/ml lysozyme. Cells were sonicated on ice as described previously. The sample was centrifuged at 14,000 x *g* for 2 hours at 4°C and the supernatant was discarded. The pellet was resuspended in 20 ml of lysis buffer + 1% n-Dodecyl-B-D-maltoside (DDM / Gold Biosciences) + 20 mM imidazole and incubated for 2 hours at 4°C on an end-over-end rotor to extract membrane proteins. The sample was then centrifuged at 14,000 x *g* for 1 hour at 4°C to separate cell debris from extracted membrane proteins. The supernatant was run over 5 ml of cobalt resin (Thermo Fisher Scientific) equilibrated with lysis buffer + 0.05% DDM + 20 mM imidazole in a gravity flow column. The column was washed with 100 ml of 20 mM sodium phosphate, 500 mM NaCl, pH 7.0 + 0.05% DDM + 20 mM imidazole. The protein was eluted in 20 mM sodium phosphate 500 mM NaCl pH 7.4 + 500 mM imidazole + 0.05% DDM and dialyzed with a 10 kDa MWCO dialysis tubing against 20 mM sodium phosphate 150 mM NaCl pH 7.0 overnight at 4°C. For gel filtration the sample was concentrated to 500 µl and run over a Superdex 200 column equilibrated with 20 mM sodium phosphate pH 7.0, 500 mM NaCl + 0.05% DDM. Sample quality was assessed by SDS-PAGE followed by Colloidal Coomassie staining or via immunoblotting using SDS-PAGE separation followed by transfer to a PVDF membrane using the Bio-Rad Trans-Blot Turbo transfer system, blocking with Odyssey blocking buffer (Bio-Rad) for 1 hour shaking at room temperature, then probing with α-6x-histidine antibody (Rabbit polyclonal, Thermo Scientific Fisher: PA1-983B)at 1:1000 in TBST overnight at 4°C shaking, followed by 3x washes for 15 minutes each with TBST, followed by secondary detection with Goat-anti-rabbit-680RD in TBST + 0.01% SDS for one hour at room temperature, followed by 3x washes with TBST, and resuspension of the membrane in 1X TBS. The blot was then imaged using an Odyssey Imager. Proteins were stored in 50% glycerol at -20°C until needed.

### CD-NTase enzymatic assays and detection of 2′,3′-cGAMP via ELISA

1 µg of recombinant CD-NTases / CdnB enzymes (from *Bacillus thuringiensis* 407 or *Clostridium botulinum*) or MBP-hcGAS were incubated at 37°C with 250 nM ATP and 250 nM GTP in 10 mM Tris-HCl pH 7.4, 25 mM KCl, 20 mM MgCl_2_, 1 mM MnCl_2_, 1 mM DTT in 200 µl total volume in in triplicate in a 96 well plate sealed with parafilm. Samples were then split in half and one set was treated with VacV poxin for 2 hours at 37°C. The samples were then analyzed using the 2′,3′-cGAMP ELISA kit (Arbor Assays) via the manufacturer’s instructions. The data was analyzed in GraphPad Prism using 4PLC analysis to interpolate the standard curve and unknowns and cross-validated using the manufacturers template. Synthetic cyclic oligonucleotides were used at 100 nM to determine cross-reactivity and agreed with the manufacturer’s observations. All synthetic oligonucleotides were obtained from BioLog/Axxora except for 2′,2′-cGAMP, which was obtained from Invivogen. Experiments were repeated 3 times with similar results, n = 2 for samples including VacV poxin treatment.

### Strain Construction in *Bacillus subtilis*

Plasmids containing indicated defense systems were transformed into *Bacillus subtilis* PY79 (hereafter referred to as Bsu) using an integrative vector (pLOCO2) with homology to the 3′ and 5′ ends of the *amyE* gene as described in Doron et al., 2018 and selected on LB + 100 µg / ml Spectinomycin (Spec100) overnight at 37° C. Successful transformants were grown overnight in LB Spec100 and frozen as 15% glycerol stocks for further use. Genomic DNA was isolated from successful transformants using the Qiagen DNAeasy blood and tissue kit and were confirmed for successful integration using PCR.

### Phage defense experiments on solid agar

Bsu strains were grown overnight at 37°C in LB supplemented with 100 µg/ml spectinomycin. Overnight cultures were diluted 1:1000 in Bsu Infection media (LB + 10 mM MgCl_2_, 10 mM CaCl_2_, and 0.1 mM MnCl_2_) and grown in 1 ml volume in a 12 ml culture tube (to allow sufficient aeration), at 30°C at 200 RPM until mid-log phase (OD600 = ∼ 0.5). 400 µl of Bsu was mixed with 4.5 ml of 0.35% LB top agar (containing 10 mM MgCl_2_, 10 mM CaCl_2_, and 0.1 mM MnCl_2_ stored at 55°C until use), inverted, and overlaid on an LB agar plate and allowed to cool for 30 minutes to create the Bsu-containing top-agar. The indicated phages were then serially-diluted, spot-plated, and plates grown overnight at 30°C. Plaques were imaged and counted the next morning and quantified as the efficiency of plaquing (PFU/ml) and displayed as histograms in Graphpad Prism.

### Phage growth curve experiments in liquid culture and viral titer measurements

Bsu strains were grown overnight at 37°C in LB + Spec100. Strains were back-diluted 1:100 in 25 ml infection medium (LB + 10 mM CaCl_2_, 10 mM MgCl_2_, 0.1 mM MnCl_2_) and grown at 30°C at 200 RPM until an OD_600_ of 0.4 was reached. Then, SPP1 phage at the indicated multiplicity of infection (MOI) was added. Following phage addition, 1 ml of sample was taken every 30 minutes, spun at 6000 x *g*, the supernatant was taken and 1 µl of chloroform was added to kill any remaining bacteria, preserving only viable phage particles. Phage samples from infection were then serially-diluted and spot plated on Bsu PY79 top agar as described above to determine the change in titers for strains containing BtCBASS or BtCBASS^CD^ strain. An increase in titers over the time course was interpreted as successful viral replication and propagation, whereas a flatline was interpreted as abortive infection due to the lack of viral replication.

### Phage amplification and isolation experiments

*Bacillus subtilis* phage SPP1 was a kind gift from Daniel Kearns. SPβ and phi29 were obtained from the DSMZ stock collection. Phages were plaque purified on Bsu PY79 in LB infection media and mixed with BsuPY79 and amplified on agar plates. Phage was scraped from the plates using a cell scraper and 100 µl of chloroform was added to lyse remaining bacteria, mixed, and allowed to settle. Phage stocks were stored at 4°C. Isolation of wild *Bacillus subtilis* phage phiB002 was performed as follows: soil from outside of the University of Colorado Jennie Smoly Caruthers Biotechnology Building was collected, mixed with LB for one hour, and then *B. subtilis* PY79 at log phase (OD 600= 0.4) was added to the mixture and allowed to grow for 3 hours. The mixture was then filter-sterilized and chloroform-treated, and single plaques were isolated on *Bsu* PY79. The phage phiB002 was isolated, plaque-purified three times, and then stored as described above. All phage stocks were validated using whole genome sequencing (SeqCoast) and were assembled to the reference genomes in Geneious Prime. phiB002 was identified as related to the myophage SBSphiJ/Grass.

### Microscale Thermophoresis (MST)

MST was performed on a NanoTemper Monolith (University of Colorado Boulder, Biochemistry Shared Instruments pool). BtCap14_6xhis_ was diluted from a stock solution into 10 mM HEPES, 1M KCl, pH 7.2, + 0.5% DDM and was labeled using a His-Tag Labeling Kit RED-tris-NTA 2^nd^ Generation Kit (NanoTemper) via the manufacturers protocol in 1.7 ml low-adhesion tubes. The sample was then spun at 16,000 x *g* after labeling to remove aggregates. The samples were then prepared with the required serial dilutions of cyclic dinucleotides in PCR tubes and were immediately transferred to Nanotemper glass capillaries and analyzed. Replicates were analyzed using the MO-Control software to obtain fraction bound, and the resulting data was plotted using GraphPad Prism. Data was fit the Hill Equation and was similar to the value obtained using the MO-Control software.

### Planar Lipid Bilayer experiments

Electrophysiology experiments were performed on an Orbit Mini (Nanion Technologies) horizontal lipid bilayer system and data was collected using Elements Data Reader 3.0. Briefly, using the manufacturers recommendations: 100 µM MECA chips (Ionera) were bathed in recording solution (150 µl) at 2 nA gain, 1.25 kHz sampling rate at + 10 mV. 1,2-diphytanoyl-sn-glycero-3-phosphocholine (DphpC) from Avanti Polar Lipids was dissolved at 10 mg/ml in n-octane (Sigma, electronics grade) and saved at -20°C until use. Membranes were painted using a paintbrush and Teflon was used to remove excess lipid until the capacitance was between 15-30 pF (100 µM MECA chip) or 30-60 pF (150 µM MECA chip) as per the manufacturer’s recommendation. Membrane integrity was monitored for 5 minutes to assess stability. Approximately 10 ng of BtCap14 was added per aperture and each membrane was monitored for fusion spikes to indicate insertion of proteins with the membrane, the fusion pulse option was used to enhance fusion at + 40 mV. 1 µM of 2′,3′-cGAMP was added in proximity to the apertures and mixed carefully using a pipette. The current trace was monitored for openings at which point either single channel data was collected at the indicated voltage or current/voltage relationships were determined. Data was analyzed using Elements Data Analyzer 3.0. All experiments were conducted at +40 mV unless otherwise indicated. For current/voltage relationships the voltage was clamped at -100 mV and activity was recorded for 10 seconds at 20 mV increments systematically with a return to 0 mV between steps. For ion substitution measurements experiments were conducted as described above, but for K-Gluconate 10 mM KCl was included in the solution to account for the use of Ag/AgCl_2_ electrodes in the experimental setup. For Gadolinium inhibition, 1 mM of gadolinium chloride was added at the end of an experiment in which open channels were present.

### Chimera10 DNA Degradation assays

Linear PCR product was amplified by1 µg of Chimera10 was incubated with 1 µM of the indicated cyclic dinucleotide with 500 ng of linear PCR-amplified DNA in a final buffer of 10 mM Tris-HCl pH 7.4, 25 mM KCl, 20 mM MgCl_2_, 1 mM DTT for 1 hour at 37°C. DNA degradation was visualized on 1% agarose gels stained with SYBR-Safe.

### DNAse alert assay for Chimera10 nuclease activity

DNAse Alert was purchased from IDT. One tube was resuspended in 5 ml of reaction buffer as described in the previous section for linear PCR product degradation. In a 96 well plate, samples were prepared in 100 µl buffer + substrate with the indicated cyclic dinucleotide stock (10 µl volume was added total) + 3 µg of Chimera10. The reaction was monitored at an excitation / emission of 360 / 460 nm in a TECAN Spark plate reader at 37°C. Data was plotted in Graphpad Prism.

### Alphafold modeling of BtCap14

The BtCap14 monomer and multimer models were generated using Alphafold 2.0 (*63*). Atomic modeling was performed in UCSF ChimeraX (*64*).

### Domain analysis of BtCap14

The BtCap14 amino acid sequence was analyzed using TMHMM (https://dtu.biolib.com/DeepTMHMM) server (*65*) and MemSat Software on the PSIPRED server, (http://bioinf.cs.ucl.ac.uk/psipred/) (*66*).

### Plasmids, Strains, Chemicals and Reagents

For a full list of plasmids, strains, and phages used in this study please refer to supplementary tables 01 – 03, respectively, which are included as a supplementary excel file.

### Accension numbers

The ascension numbers for the indicated genes are: DaCdnB, WP_031517737.1 ; BtCdnB, EEM25276.1; CbCdnB, WP_053342861.1; BtCap14, WP_098368803.1, CbCap14, WP_053342862.1, GbCap5; WP_013400843.1.

## Supplementary Data Tables

**Table S1**. Plasmids used in this study

**Table S2**. Bacterial strains used in this study.

**Table S3**. Phages used in this study.

## Supplementary Figures

**Figure S1.**
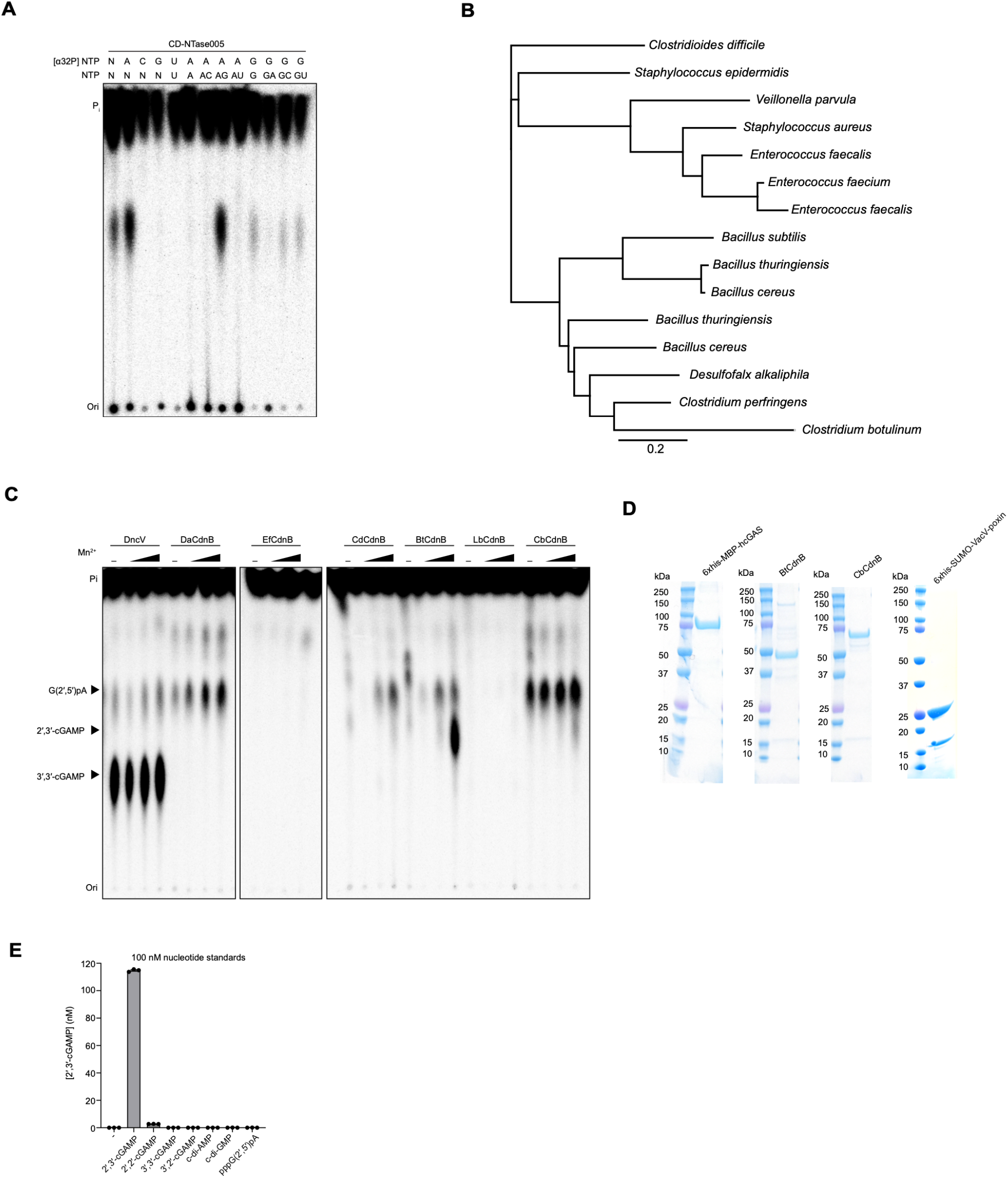
Bacterial CdnB enzymes and their products. **(A)** PEI-cellulose TLC analysis of CD-NTase reaction products produced when CD-NTase005 was incubated with the indicated α^32^P-radiolabelled and unlabeled NTPs. **(B)** Phylogram of CD-NTase005 homologues. The key represents substitutions per site. **(C)** PEI-cellulose TLC analysis of enzyme reaction products as in Fig. 1A with the indicated CD-NTase005 homologue and Mn^2+^ added at either 0, 0.01, 0.1, or 1 mM. The approximate migration of cGAMP and linear intermediate are noted. **(D)** Coomassie stained SDS-PAGE gel of 6xhis-MBP-hcGAS, CdnB enzymes, and 6xhis-SUMO-VacV poxin enzyme used in this study. **(E)** 2′,3′-cGAMP ELISA quantification of 100 nM solutions of synthetic cyclic dinucleotides to determine cross-reactivity.

**Figure S2.**
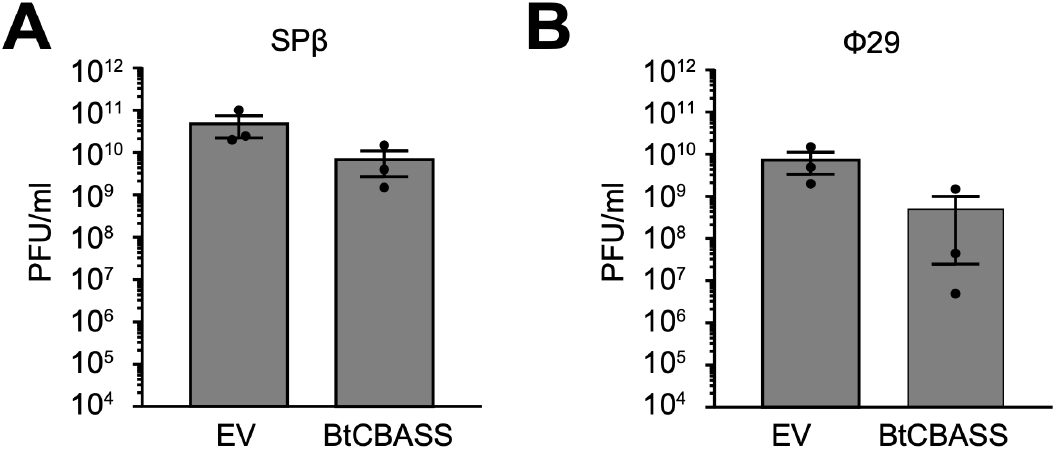
*B. subtilis* expressing BtCBASS and challenged with SPβ or Φ29. Efficiency of plating of the indicated phage infecting *B. subtilis* expressing empty vector (EV) or BtCBASS. Data represent mean ± SEM for n = 3 biological replicates.

**Figure S3.**
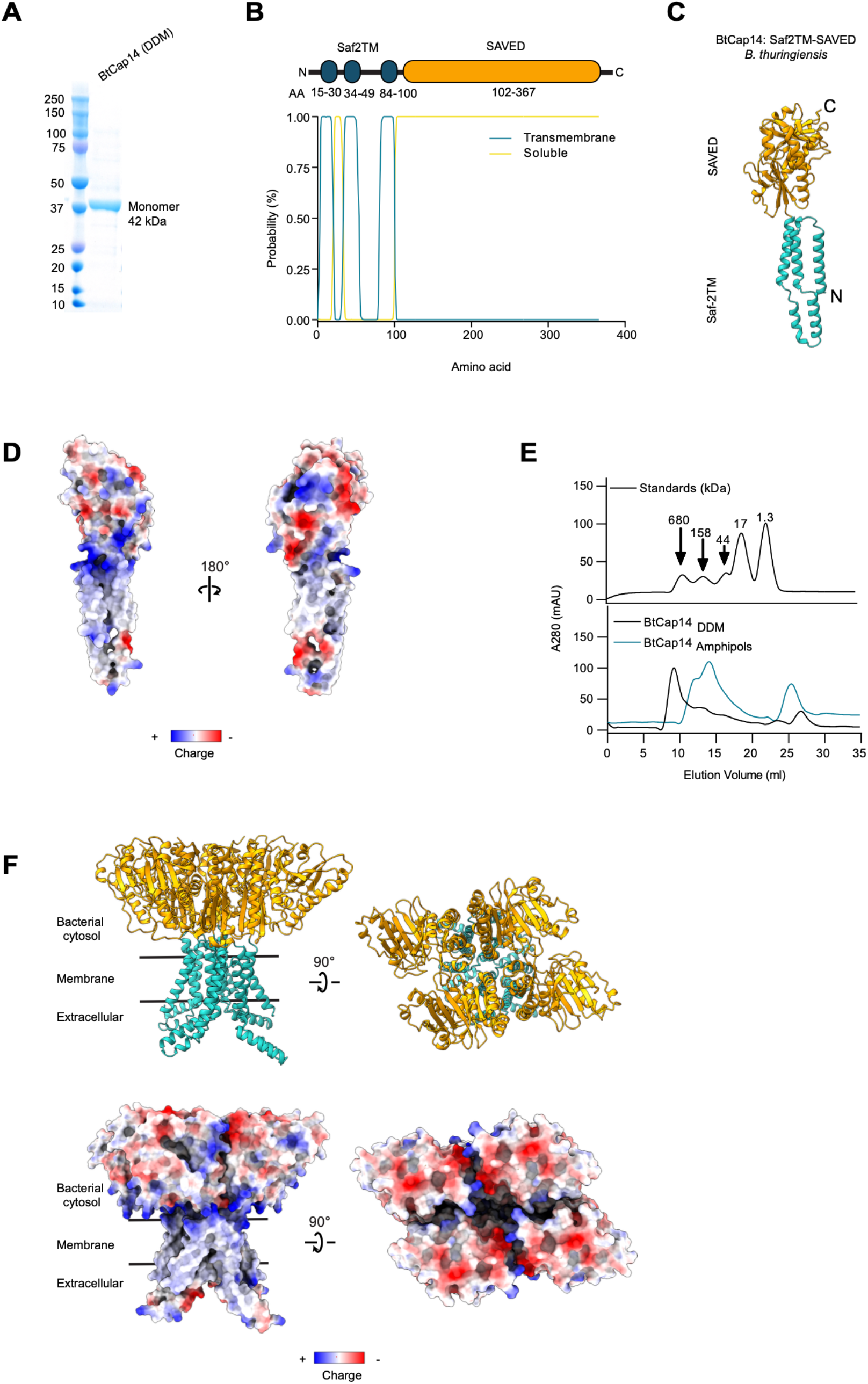
Purification, analysis, and modeling of BtCap14. **(A)** Coomassie-stained SDS-PAGE gel of BtCap14_6xhis_ purified in n-Dodecyl-B-D-Maltoside (DDM) after IMAC and gel-filtration. **(B)** Domain annotation of BtCap14 using TMHMM software. Shown is a probability plot of the amino acids predicted to be transmembrane or soluble with a cartoon diagram above. **(C)** Predicted structure of monomeric BtCap14. The Saf-2TM (blue) and SAVED (goldenrod) domains are at the N and C termini, respectively. **(D**) Model from (C) colored according to electrostatic charge. **(E)** Gel filtration analysis of BtCap14_6xhis_ in DDM micelles or amphipols on a Superdex 200 10/30 column in 20 mM sodium phosphate pH 7.2, 150 mM KCl, + 0.05% DDM. The top panel represents protein standards run under identical conditions. **(F)** Predicted structure of the BtCap14 homotetramer using multimer modeling. The tetramer, colored as in (C) is arranged in a lipid bilayer according to analysis in (B). Shown below is the same model colored according to electrostatics.

**Figure S4.**
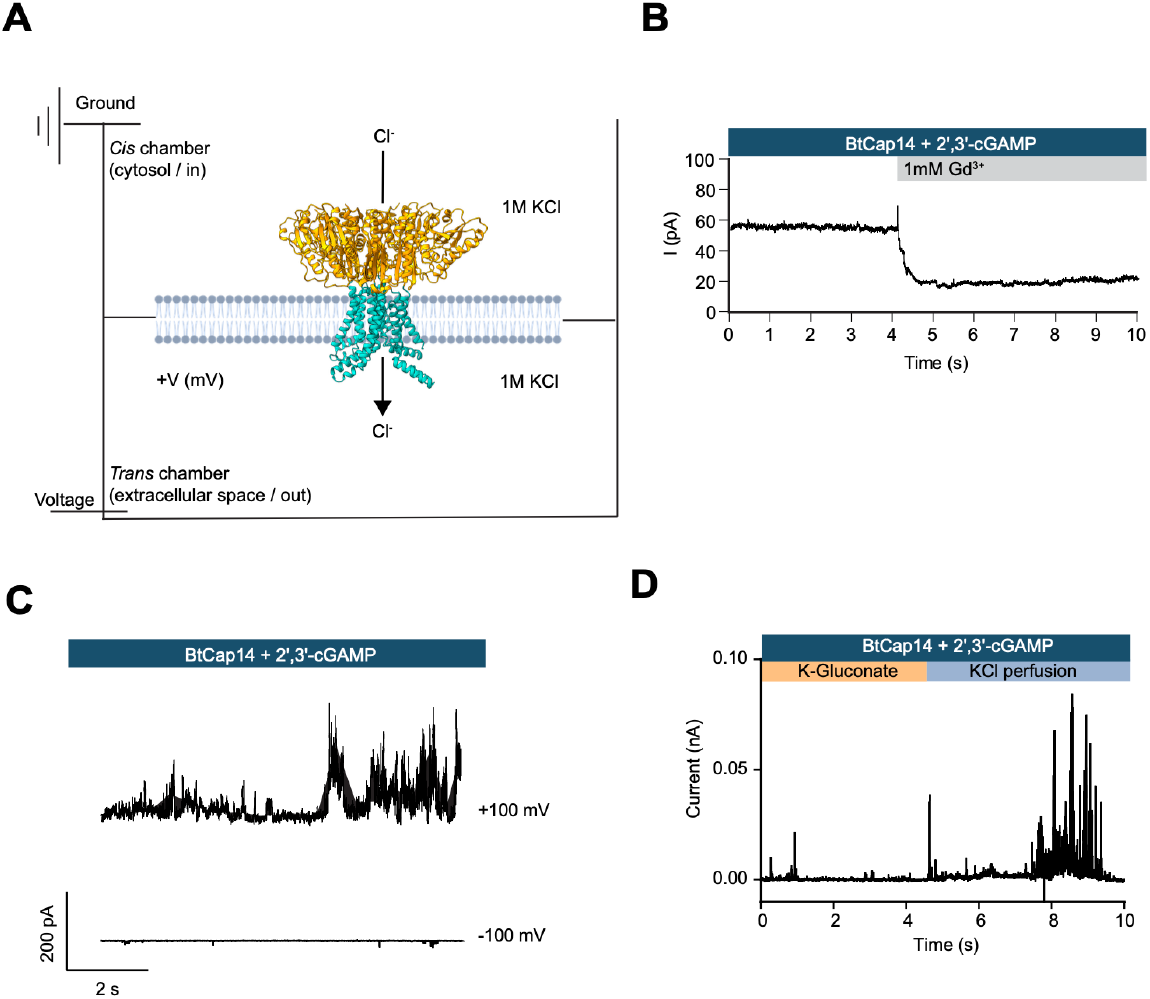
Electrophysiological analysis of BtCap14. **(A)** Schematic of planar lipid bilayer set up and electrophysiological convention used in this study. Current at positive holding potentials (+V_m_) and analysis of alternative anions demonstrates chloride transport from *cis* to *trans*, consistent with BtCap14-mediated Cl^-^ export from the cell in the presence of 2′,3′-cGAMP. **(B)** Inhibition of BtCap14 channels by gadolinium (Gd3^+^) chloride in the presence of 2′,3′-cGAMP. Data are representative of n = 3 biological replicates. **(C)** Current trace of BtCap14 + 2′,3′-cGAMP at the indicated holding potential. Data are representative of n = 3 biological replicates. **(D)** Channel rescue experiment. Current trace in which BtCap14 + 2′,3′-cGAMP was reconstituted in 1 M K-Gluconate + 10 mM KCl, and then perfused with 1 M KCl on the *cis* side of the bilayer. Data are representative of n = 3 biological replicates.

**Figure S5.**
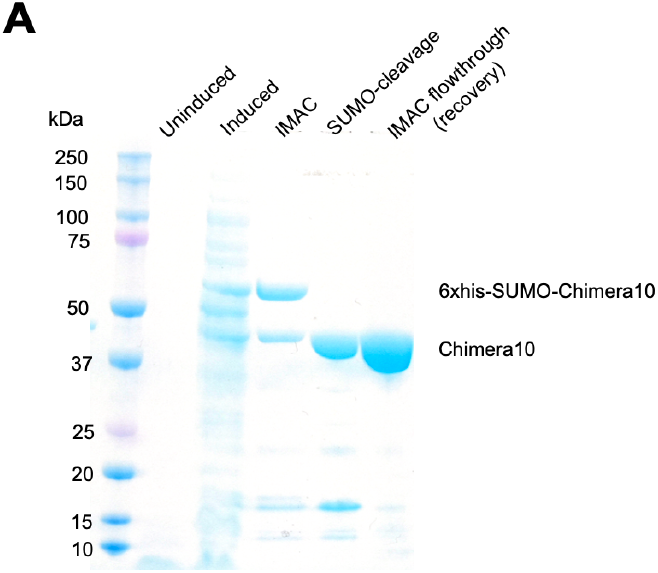
Purification of Chimera10. **(A)** Coomassie stained SDS-PAGE gel indicating expression and purification steps of 6xhis-SUMO-Chimera10 (Final monomeric molecular weight after tag cleavage: 45 kDa).

